# Protein expression of short-chain dehydrogenases/reductases and their inducibility by flubendazole in *Haemonchus contortus*

**DOI:** 10.64898/2026.08.07.743491

**Authors:** Michaela Šadibolová, Karolína Štěrbová, Nikola Slaninová, Nicole Lübbehusen, Thomas Ruppert, Marcin Luzarowski, Petra Matoušková, Lenka Skálová, Lucie Raisová Stuchlíková

## Abstract

Short-chain dehydrogenases/reductases (SDRs) constitute a large enzyme superfamily involved in endogenous metabolism and xenobiotic biotransformation. In the parasitic nematode *Haemonchus contortus*, SDRs may catalyze the carbonyl reduction of a benzimidazole anthelmintic flubendazole (FLU), whose increased reduction is associated with FLU resistance.

This study thus investigated the constitutive expression of SDRs and their inducibility by FLU in drug-susceptible and benzimidazole-resistant strains of *H. contortus*. The expression of 23 sdr genes was analyzed by quantitative PCR, while targeted proteomic assays enabled the quantification of 15 SDR proteins. In adult nematodes, pronounced sex-dependent differences were detected at both transcript and protein levels. Resistance-associated alterations were less pronounced and were observed predominantly in males, with SDR9, SDR12, SDR15, and SDR20 displaying increased protein abundances in the resistant strain. Exposure to FLU induced only minimal transcriptional responses in juvenile stages, whereas adult nematodes exhibited marked sex- and strain-specific responses in the expression of SDRs. The strongest transcriptional effects were detected in resistant males, while significant protein-level changes following FLU treatment were observed exclusively in adults of the drug-susceptible strain. Notably, SDR9 and SDR20 combined resistance-associated expression patterns with responsiveness to FLU exposure. Taking together, the first targeted proteomic characterization of SDRs in *H. contortus* revealed several SDR isozymes with constitutive overexpression in resistant nematodes and/or inducibility by FLU, suggesting a potential role in adaptation to anthelmintic exposure.

## Introduction

*Haemonchus contortus* (family *Trichostrongylidae*) is a blood-feeding parasitic nematode that infects ruminants. Animals with haemonchosis exhibit symptoms such as anemia, bottle jaw (swelling of the lower jaw), edematous stomachs (bloating due to fluid accumulation in the abomasum), and overall emaciation. Due to reduction in growth rates, decreased milk production, reduced wool quality (in the case of sheep), lower overall productivity and high mortality, haemonchosis of food-producing animals is accompanied with considerable economic losses for their farmers [1]. Pharmacotherapy using anthelmintics represents the primary strategy for the treatment of haemonchosis. However, overuse and misuse of anthelmintics have led to the development of resistance among nematodes, making it more difficult for these drugs to treat haemonchosis effectively. Moreover, the prevalence of multidrug resistance in *H. contortus* to all classes of anthelmintics has increased worldwide [2–4]. Therefore, the mechanisms of resistance development have been intensively studied.

Concerning benzimidazole anthelmintics, changes in their target molecule, β-tubulin, have been demonstrated as the primary mechanism of nematode resistance [2]. In addition, non-target site mechanisms, such as the increased expression and activity of drug-metabolizing enzymes (DMEs), further contribute to drug resistance in *H. contortus*. It is known that adult *H. contortus* from a drug-resistant strain are able to deactivate several benzimidazole anthelmintics more efficiently in comparison with the adults from a drug-sensitive strain [5]. Flubendazole (FLU) [6], an anthelmintic drug bearing a carbonyl group, is deactivated via carbonyl reduction in *H. contortus*. Reduction of a carbonyl group is a typical phase I biotransformation reaction that increases hydrophilicity and enables subsequent conjugation, thereby facilitating drug excretion [7]. In the drug-resistant strain of *H. contortus*, FLU was reduced to a greater extent than in the drug-sensitive strain [5, 8, 9]. The comparison of FLU reduction across different developmental stages of *H. contortus* (eggs, L1/2, L3, xL3, adult males and females) confirmed that FLU was reduced in all developmental stages, with adult females being the most metabolically active [9]. Moreover, *in vivo* exposure of *H. contortus* to sublethal doses of FLU led to increased FLU reduction. Data from the inhibition study revealed that menadione was an effective inhibitor of FLU reduction in adult *H. contortus* both *in vitro* and *ex vivo*. Menadione is a well-known substrate and a competitive inhibitor of human carbonyl reductase 1, which belongs to the short-chain dehydrogenases/reductases (SDRs) superfamily [5, 10]. In view of this fact, participation of some SDR enzymes in FLU reduction in *H. contortus* can be assumed and deserves further attention.

SDRs constitute a large and functionally diverse superfamily of enzymes involved in oxidation-reduction reactions. They typically catalyze the transfer of hydrogen between substrates using nicotinamide cofactors NAD(H) or NADP(H). Structurally, SDRs share a conserved Rossmann-fold domain responsible for cofactor binding and an active site with a characteristic Tyr-Lys-Ser triad that facilitates catalysis. Although only about 15-30% of their amino acids are identical, these proteins retain a very similar overall three-dimensional structure. SDRs play crucial roles in metabolism, including steroid biosynthesis, xenobiotics detoxification, and secondary metabolite production. Their broad substrate specificity and functional versatility make them interesting targets for drug development and biotechnological applications [11–13]. In *H. contortus*, genome sequencing enabled the identification of 46 members of SDR family with SDR1, SDR3, SDR5, SDR6, SDR14, and SDR18 genes being the most expressed across all developmental stages. A transcriptomic comparison of SDRs levels between the drug-susceptible and drug-resistant strains of *H. contortus* revealed an increased expression of SDR1, SDR12, SDR13, and SDR16 in most developmental stages of the latter [14]. However, the protein expression and inducibility of SDRs in *H. contortus* have not yet been studied.

As previously reported, exposure of nematodes to sublethal doses of several anthelmintic drugs can increase the expression and activity of DMEs, mainly those catalyzing drug deactivation [15, 16]. Therefore, we investigated whether exposure of *H. contortus* to sublethal doses of FLU could induce the expression of specific SDRs. The present study was designed to measure the mRNA expression of selected SDR enzymes across all developmental stages of *H. contortus* exposed to various doses of FLU (0.01µM, 1µM and 5µM). Furthermore, we performed a targeted proteomic analysis to evaluate the constitutive protein levels of selected SDRs in adult *H. contortus*, focusing on sex-related differences and differences between the drug-susceptible (ISE) strain and the benzimidazole-resistant (IRE) strains.

## Materials and methods

### Parasites and their collection

Two isolates of *H. contortus* were used: an inbred susceptible-Edinburgh strain (ISE, MHco3) and an inbred resistant-Edinburgh strain (IRE, MHco5) [17]. Sheep, the hosts of *H. contortus*, were bred and slaughtered in agreement with the Czech slaughtering rules for farm animals and the Protection of Animals from Cruelty Act No. 246/1992, Czech Republic. The breeding facilities were accredited by the Ministry of Agriculture of the Czech Republic for experimental sheep housing (Approvement MZE-53255/2022-13143). All experimental procedures were evaluated and approved by the Ethics Committee of the Ministry of Education, Youth and Sports of the Czech Republic (Project number MSMT-20144/2023-4).

Adults of *H. contortus* were obtained using our routine procedure published previously [16, 18]. Briefly, parasite-free lambs were orally infected with 6,000 third-stage larvae (L3s) of either the ISE or IRE strain of *H. contortus*. Eggs were isolated from faeces and refined with a sucrose flotation technique followed by washing in tap water. The mixture of first-stage and second-stage larvae (L1/L2s) developed from isolated eggs incubated in tap water at 27 °C for 24 hours. L3s were produced by incubating eggs in humidified faeces at 27 °C for one week, after which they were washed in tap water and separated using sieves with different mesh sizes. To obtain exsheathed L3s (xL3s), L3s were exposed to 0.15 % (v/v) of sodium hypochlorite (NaClO) at 37 °C for 20 minutes [19]. Lambs were stunned and immediately exsanguinated seven weeks after infection. The agar method was used to obtain adult nematodes from lambs’ abomasa. Sexually dimorphic adults were sorted manually according to sexual characteristics. [20].

### Exposure of *H. contortus* to FLU

For the targeted transcriptomic analysis, 75,000 isolated eggs, 75,000 L1/L2s, 30,000 L3s, 30,000 xL3s, 10 females, and 15 males of *H. contortus* from the ISE and IRE strains were used per sample to obtain sufficient and comparable amounts of RNA [18]. Eggs and L1/L2s were exposed to 0.01 µM or 1 µM FLU (Janssen Pharmaceutica, New Brunswick, NJ, USA; pre-dissolved in dimethyl sulfoxide (DMSO). L3s, xL3s, females, and males were exposed to 0.01 µM, 1 µM FLU or 5 µM FLU, as they tolerate higher concentrations of FLU. Developmental stage-matched controls were exposed to DMSO only and were incubated under the same conditions. The final concentration of DMSO in the medium was 0.1 %. Samples from each developmental stage were prepared in four biological replicates. Adult nematodes were incubated in a RPMI 1640 medium (Sigma-Aldrich, Prague, Czech Republic) at 37 °C in a CO_2_/O_2_ incubator (5 % CO_2_) for 4 and 12 hours. Eggs and L1/L2s were incubated in tap water for 12 hours at 27 °C, whereas L3s and xL3s were incubated for 12 hours at 37 °C. After exposure, all larvae and adult nematodes remained alive, as confirmed by the motility assessment, regardless of the presence or absence of FLU. Immediately after incubation, all stages of *H. contortus* were washed with sterile tap water, placed into 1 mL of TriReagent® (Molecular Research Centre, OH, United States) and stored at −80 °C for later use.

For the targeted proteomic analysis, 10 females and 20 males from the ISE and IRE strains were incubated separately in a RPMI 1640 medium (Sigma-Aldrich, Prague, Czech Republic) with 5 µM FLU (pre-dissolved in DMSO) or DMSO only (controls) at 37 °C in a CO_2_/O_2_ incubator (5 % CO_2_) for 24 hours. The final concentration of DMSO in the medium was 0.1 %. Each adult group was prepared in four biological replicates. After exposure, all samples were homogenized in 600 µL of 8M urea in 100 mM tetraethylammonium bromide (TEAB) using a FastPrep-24 5G Homogenizer (MP Biomedicals, France) (two 40 s intervals, 6 m/s) and using three 30-second cycles of sonication (Sonoplus HD 2070, Bandelin, Berlin, Germany). Supernatant was obtained by centrifugation (12,000 *g*, 4 °C, 20 min) and stored at −80 °C until further processing. Protein concentration of supernatants was estimated using a Bradford assay (Bio-Rad, CA, USA) according to the manufacturer’s instructions.

### RNA isolation and cDNA synthesis

Total RNA was extracted using TriReagent® according to the manufacturer’s protocol. All samples were homogenized in a FastPrep-24 5G Homogenizer (MP Biomedicals, France) for four 30 s intervals with a speed of 6.5 m/s. The purity and concentration of RNA were determined spectrophotometrically using a NanoDrop ND-1000 UV-Vis Spectrophotometer (Thermo Fisher Scientific, MA, USA) and analyzed by an Agilent 2100 Bioanalyzer on RNA Nano Chips (Agilent Technologies, CA, USA). To avoid DNA contamination, 4 µg of RNA from each sample were treated with DNase I (New England Biolabs, UK) and diluted to a concentration of 0.1 µg/µL. Complementary DNA (cDNA) was synthesized from 0.5 µg of RNA in 20 µL reactions using random hexamer primers and Protoscript II Reverse Transcriptase (New England Biolabs, UK) following the manufacturer’s protocol. First-strand DNA was diluted 10x and stored at −20 °C until further analysis.

### Quantitative PCR (qPCR)

Eight replicates of each sample (four biological replicates, each in two technical replicates) were analyzed in 384-well plates. The final mastermix (8 µL per well) was prepared from 5 ng of cDNA, qPCR Xceed SG 1 step 2x Mix Lo-ROX (IAB, Czech Republic), and both forward and reverse primers at a final concentration of 100 nM. qPCR assay was performed in a 384-Well PCR Thermal Cycler and a QuantStudio^TM^ 6 Flex Real-Time PCR System (Applied Biosystems, CA, USA) with SYBR Green I detection. The qPCR run protocol started with an initial denaturation step (95 °C for 2 min), followed by 40 cycles of two-step amplification (95 °C for 15 s, 60 °C for 20 s), with fluorescence measured at the end of each cycle. The melting curve protocol with a heating rate of 0.5 °C every 30 s starting from 60 °C to 95 °C was used to investigate the specificity of the qPCR reaction. Gene-specific amplification was confirmed by observing a single peak in each sample. For normalization of the qPCR assay, a combination of two reference genes, glyceraldehyde-3P-dehydrogenase (*gpd*) and nuclear cap-binding protein subunit 2-like (*ncbp*), was used as recommended by Lecová et al. [21]. Primer sequences for quantification of SDR genes, designed and verified in our previous study [14], were synthesized by Generi Biotech, Czech Republic.

### Statistical analysis of mRNA expression data

Differences in the constitutive mRNA expression between sample groups were assessed using a pairwise linear regression analysis followed by Benjamini-Hochberg correction for multiple comparisons in the same way as the protein expression data (see *Targeted proteomics data and statistical analysis*). The data from FLU incubations are expressed as the mean ± S.D. of 4 biological replicates. Relative mRNA expression was calculated using the ΔΔCt method [22]. Normal distribution of the expression data was assessed using the Shapiro-Wilk normality test (n=4, *p*<0.05). Statistical significance of FLU-treated samples from the sensitive and resistant strains was evaluated using two-way ANOVA, with the control group set to 1, followed by Dunnett’s *post hoc* test. These results were processed in GraphPad Prism 10.

### Protein extraction and sample preparation for targeted LC-MS measurements

Three hundred µg of protein was precipitated using chloroform-methanol precipitation [23]. Protein pellets were resuspended in 20 µL of 8 M urea (in 100 mM TEAB) and protein concentration was estimated again using the Bradford assay. Afterwards, 100 µg of protein was reduced and alkylated with 10 mM tris(2-carboxyethyl)phosphine hydrochloride (TCEP) and 40 mM 2-chloroacetamide (CAA), respectively, for 30 min at room temperature. Samples were diluted with 50 mM TEAB to a final urea concentration of 1.6 M and digested with MS-grade trypsin at 1:50 (w/w) enzyme-to-protein ratio overnight at 37 °C. After digestion, samples were acidified with trifluoracetic acid (TFA) to reach pH < 2. Samples were desalted using Sep-Pak tC18 100 mg cartridges (Waters, MA, USA). Peptides were eluted during two consecutive washes with 50 % acetonitrile (ACN) with 0.1 % TFA and 80 % ACN with 0.1 % TFA, respectively. Samples were vacuum-dried and stored at −20 °C until further LC-MS analysis.

### Targeted proteomic assay development

Twenty SDR proteins were selected based on our previous work [14]. Their UniProt and WormBase accessions along with their protein sequences are listed in Additional file 1. For the development of the targeted proteomic analysis, 2-3 peptides for each protein were selected based on an in-house acquired shotgun proteomic dataset fulfilling the following criteria: proteotypicity, sequence length between 7-22 amino acids, no methionine in the sequence, no missed cleavages or ragged ends, and a good intensity of their MS signal. For SDR proteins identified with only one peptide or not identified at all in the shotgun dataset, 2-3 peptides determined by the *in silico* trypsin digestion were selected, taking into account the abovementioned criteria. In the case of a significant sequence overlap of some SDR proteins, proteotypicity was the major decisive factor for peptide selection. Proteotypicity of selected peptides was checked against *H. contortus* proteome (UP000025227, accessed from UniProt in February 2023) using an in-house R script.

Stable isotope labeled (SIL) peptides containing heavy lysine (^13^C_6_, ^15^N_2_) or heavy arginine (^13^C_6_, ^15^N_4_) and carbamidomethyl-modified cysteines were obtained from JPT Peptide Technologies (Berlin, Germany). Stock solutions (100 pmol/µL) were prepared in 20 % ACN with 0.1 % TFA with an addition of BSA (5 fmol/µL).

Digested samples of both strains and sexes of *H. contortus* were used as matrices to account for potential sample-dependent matrix effects and interferences. Firstly, 1 pmol of SIL peptides together with 5 µg of *H. contortus* digested protein and 1 pmol of indexed retention time (iRT) peptides were injected into a QExactive HF Orbitrap Mass Spectrometer (Thermo Fischer Scientific, Bremen, Germany) to select optimal transitions. A mixture of 10 synthetic reference iRT peptides was used for retention time calibration across chromatographic runs [24]. MS data were imported and further processed in Skyline (version 23.1.0.455) [25]. In addition, SIL peptides were also injected to a QTrap 5500 Triple Quadrupole Mass Analyzer (Sciex, Darmstadt, Germany) to compare the LC-MS behavior between the two systems. Subsequently, the best performing peptides and transitions were selected based on their intensity, retention time stability, lack of interfering signal, and detectability of endogenous counterparts. In addition, the mode of targeted analysis, particularly multiple reaction monitoring (MRM) versus parallel reaction monitoring (PRM), for individual peptides was selected based on superior LC-MS behavior between the two tested systems. This way, 15 SDRs could be reliably quantified.

For the final analysis, 1-5 transitions per peptide and 1-3 peptides per protein were used, with 21 peptides selected for a PRM analysis and 8 peptides for a MRM analysis (Additional file 2). The concentration of corresponding SIL peptides was also adjusted in their final master mix so that the light-to-heavy (L/H) peptide ratio would stay in the range of 1:4 – 4:1 (L/H). Indexed retention time was calculated using the iRT peptide standards [24], and the scheduling window was set to 5 min.

### LC-PRM and LC-MRM analyses

Nanoflow LC-PRM analysis was performed on a Dionex UltiMate 3000 RSLCnano HPLC system coupled to a QExactive HF Orbitrap Mass Spectrometer (both Thermo Fisher Scientific, Bremen, Germany). Peptide separation was achieved on an in-house packed C18 column (75 µm × 230 mm, ReproSil-Pur 120 C18-AQ, 1.9 µm particle size; Dr. Maisch, Germany). Solvent A consisted of 0.1 % formic acid in 1 % ACN (v/v) and solvent B consisted of 0.1 % formic acid in 89.8 % ACN (v/v). Samples were dissolved in 0.1% TFA to reach the protein concentration of 1 µg/µL. The final samples also contained 1 pmol of iRTs and a concentration-adjusted master mix of SIL peptides. Five µg of digested protein was injected onto the column heated to 40 °C and loaded for 20 min at a flow rate of 550 nL/min and 3 % B. Peptides were eluted during a two-step linear gradient increasing from 3% B to 37 % B over 60 minutes and further to 57.8 % B over 8 minutes at a reduced flow rate of 300 nL/min, followed by washing and reconditioning of the column to 3% B. Nanospray ionization source was operated using the following settings: spray voltage of 2.5 kV, capillary temperature of 250 °C, probe heater temperature of 350 °C, and a S-lens RF level of 60. A Skyline-exported inclusion list (Additional file 3) was provided, and corresponding fragmentation spectra were obtained using normalized collision energy (NCE) of 27 with a resolution of 30,000, an AGC target of 2e5, a maximum injection time of 50 ms, and an isolation window of 0.8 *m/z*.

Nanoflow LC-MRM analysis was carried out using a Waters nanoACQUITY UPLC System (Waters, Milford, MA, USA) coupled to a QTrap 5500 Triple Quadrupole Mass Analyzer via a NanoSpray III Source. Uncoated precut emitters (SilicaTip, i.d. 20 μm, tip i.d. 10 μm; New Objectives) and a voltage of approximately 2.6 kV were used for ESI. Five µg of digested protein (prepared in the same way as for the PRM analyses) was injected and trapped on a trapping column (Symmetry C18, 180 μm × 20 mm, 5 μm particle size, 100 Å pore size; Waters) for 7 minutes at a flow rate of 10 μL/min with 99.4 % of solvent A. Peptide separation was achieved on a C18 analytical column (M-Class Peptide BEH C18, 75 μm × 250 mm, 1.7 μm particle size, 130 Å pore size; Waters) controlled by a binary pump system delivering solvent A: 0.1 % formic acid in 1 % ACN (v/v) and solvent B: 0.1 % formic acid in 89.8 % ACN (v/v) at a flow rate of 300 nL/min. The column was maintained at 60 °C and peptides were eluted during a two-step linear gradient increasing from 3 % B to 37 % B over 37 minutes with a further increase to 57.8 % B over additional 8 minutes, followed by washing and reconditioning of the column to 3 % B. MRM measurements were performed with target scan time of 1.8 sec; collision energies were calculated using empirical equations from the Skyline software (Additional file 4).

### Targeted proteomics data and statistical analysis

Acquired LC-MS data were imported into Skyline, peak boundaries and individual transitions were manually inspected and, if needed, adjusted to ensure correct peak assignment and quantification. All light and heavy peptides had a dot-product (dotp) correlation score of > 0.83, with the vast majority > 0.90, and at least 6 points per peak. Interfering transitions were removed from the final quantification. A list of raw light and heavy peak areas was exported and subsequently analyzed in R. MRM and PRM results were processed independently. Firstly, the L/H ratio of the total peak areas was calculated for each individual measurement, followed by the calculation of the mean ratio for each peptide measured in 3 technical replicates. Subsequently, mean protein ratios were calculated by averaging across corresponding peptides, and these ratios were sum-normalized to account for sample loading variability. Finally, the sum-normalized protein ratios from both acquisition methods were combined, and the final mean ratios were computed for proteins measured using both acquisition methods (6 SDR proteins in total). These ratios were log2-transformed for downstream analysis. Pairwise group comparisons were performed using protein-wise linear regression models, with the slope of linear regression corresponding to the fold change between the tested conditions. The respective standard error of linear regression was used to calculate the 95 % confidence intervals, and the computed *p*-values were adjusted using the Benjamini-Hochberg procedure.

## Results

### Comparison of the constitutive mRNA and protein expression of SDRs in females and males of *H. contortus*

First, we examined the differences in the constitutive mRNA and protein expression of individual SDRs between males and females from the ISE and IRE strains. The expression of several investigated SDRs differed between sexes in at least one strain at both molecular levels. At the transcript level, 15 out of 23 analyzed SDRs exhibited significantly different levels, with majority of these differences observed jointly for both strains (*sdr2*, *sdr4*, *sdr5*, *sdr7*, *sdr13*, *sdr14*, *sdr16*, *sdr17*, *sdr18*, and *sdr22*) (Figure 1A). Moreover, the sex-related differences in the mRNA expression were further pronounced in the IRE strain, with 4 additional SDRs (sdr6, sdr8, sdr11, and sdr23) significantly dysregulated between sexes. Interestingly, SDR transcript levels were overall higher in females than in males in both tested strains.

**Figure 1.**
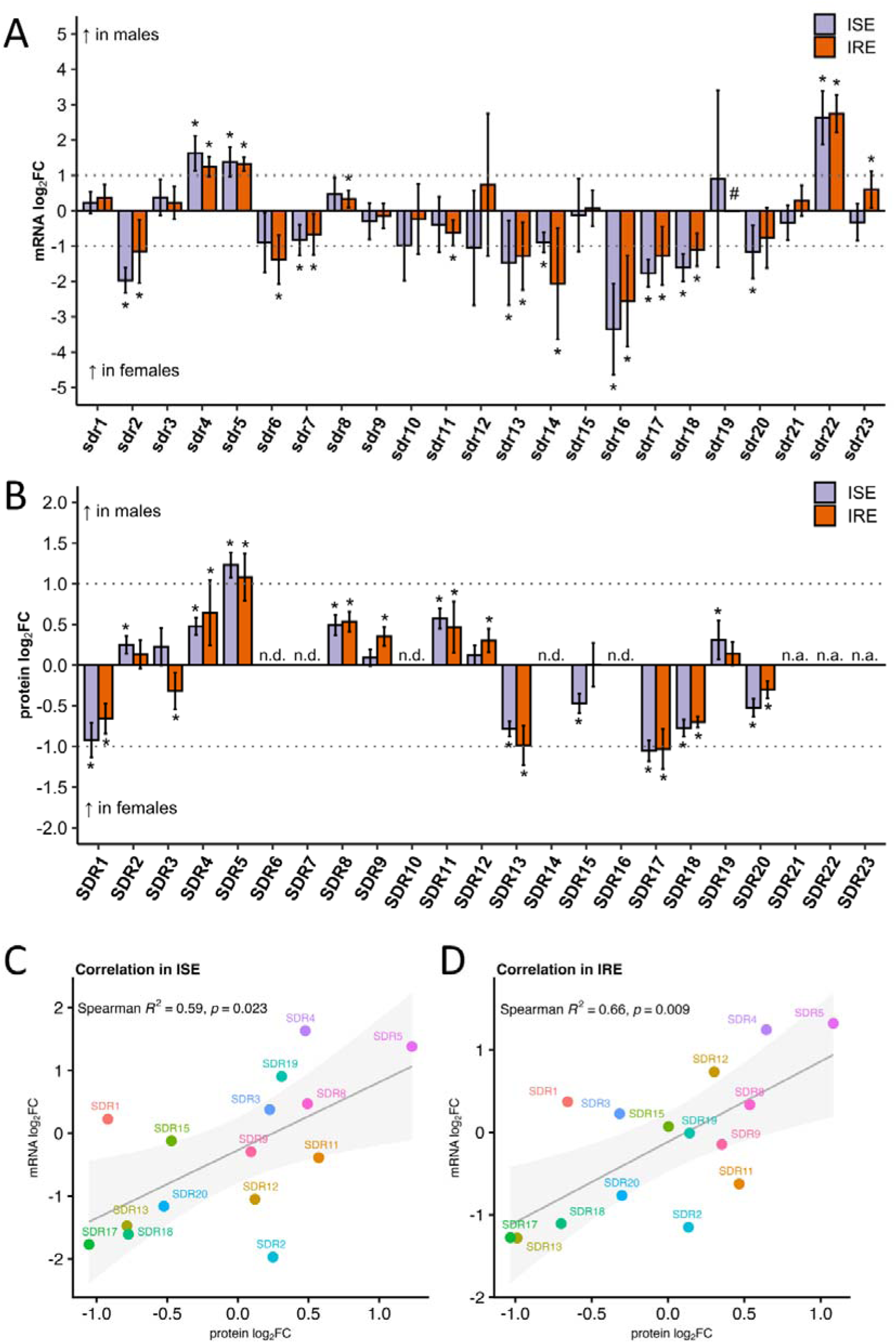
Relative comparison of the constitutive mRNA and protein expression of SDRs between males and females of *H. contortus*. Relative mRNA (A) and protein (B) expression of 23 and 15 SDRs, respectively, are displayed as log_2_ fold changes (FC) in males compared to females from the drug-sensitive (ISE) and benzimidazole-resistant (IRE) strains. mRNA data (n = 2-4) were calculated using the ΔΔCt method and normalized to the geometric mean of the two reference genes (*gpd*, *ncbp*). Protein abundances were determined by targeted MRM/PRM analysis (n = 4). For each peptide, signal intensity of the endogenous light peptide was normalized to that of the corresponding stable isotope labelled heavy peptide. Protein-level values were then calculated from the normalized peptide light-to-heavy ratios. Pairwise group differences were assessed using linear regression models, and statistical significance was determined using Benjamini–Hochberg–adjusted p-values. Error bars represent 95% confidence intervals; asterisks indicate statistically significant differences after multiple testing correction (adjusted *p* < 0.05). Horizontal dotted lines indicate log_2_FC of ± 1. (C, D) Relationship between mRNA and protein abundance of 15 SDRs assessed using linear regression. Data were derived from (A) and (B). The regression line is shown with its associated 95% confidence interval; Spearman’s rank correlation coefficients and the corresponding *p* values are indicated. Abbreviations/symbols: #, 95% confidence interval = ± 11.42; n.d., not detected; n.a., not analyzed

Using targeted proteomic investigation, we were able to robustly quantify 15 SDR proteins. There were two main limitations of this approach, in addition to the criteria for peptide selection described in the *Methods* section: (i) overlapping peptide sequences with other *H. contortus* or *O. aries* proteins (SDR6, SDR14, SDR16) (Additional file 5), and (ii) presumably low abundance of endogenous proteins (SDR7, SDR10). In addition, protein levels of SDR21, SDR22, and SDR23 were not considered at the time of instrument availability.

At the protein level, virtually all 15 measured SDR isozymes exhibited significant sex-related differences in expression in at least one tested strain (Figure 1B). As observed for the mRNA expression, most of these sex-associated differences in the protein abundance were shared between the two strains (SDR1, SDR4, SDR5, SDR8, SDR11, SDR13, SDR17, SDR18, and SDR20). In contrast to the mRNA expression, however, overall protein expression changes were distributed similarly between sexes, and additional strain-specific differences in the abundance of other SDRs were observed also equally in both strains. Specifically, in ISE adults, SDR15 was significantly upregulated in females, while SDR2 and SDR19 were elevated in males. On the other hand, in IRE adults, SDR3 was more abundant in females, and SDR9 and SDR12 in males.

Importantly, the altered mRNA expression of *sdr4*, *sdr5*, *sdr8* (in IRE), *sdr13*, *sdr17*, *sdr18*, and *sdr20* (in ISE) was consistently reflected also at the protein level, and these enzymes contributed strongly to the overall positive correlation between the transcript and protein levels among the investigated SDRs (Figure 1C, D).

### Resistance-associated changes in the protein expression of SDRs in females and males of ***H. contortus***

The comparison of the constitutive protein expression of the 15 analyzed SDRs between adults from the ISE and IRE strains is presented in Figure 2. The resistance-related changes were less pronounced compared to the sex-associated changes reported here. Nevertheless, 7 SDRs were altered in adults from the IRE strain, and, interestingly, most of this dysregulation was observed in males. Specifically, SDR2 (−10 %) and SDR5 (−19 %) were downregulated, while SDR9 (+22 %), SDR12 (+9 %), SDR15 (+48 %), and SDR20 (+20 %) were found upregulated in resistant males. In resistant females, only SDR4 (−32 %) was found significantly downregulated.

**Figure 2.**
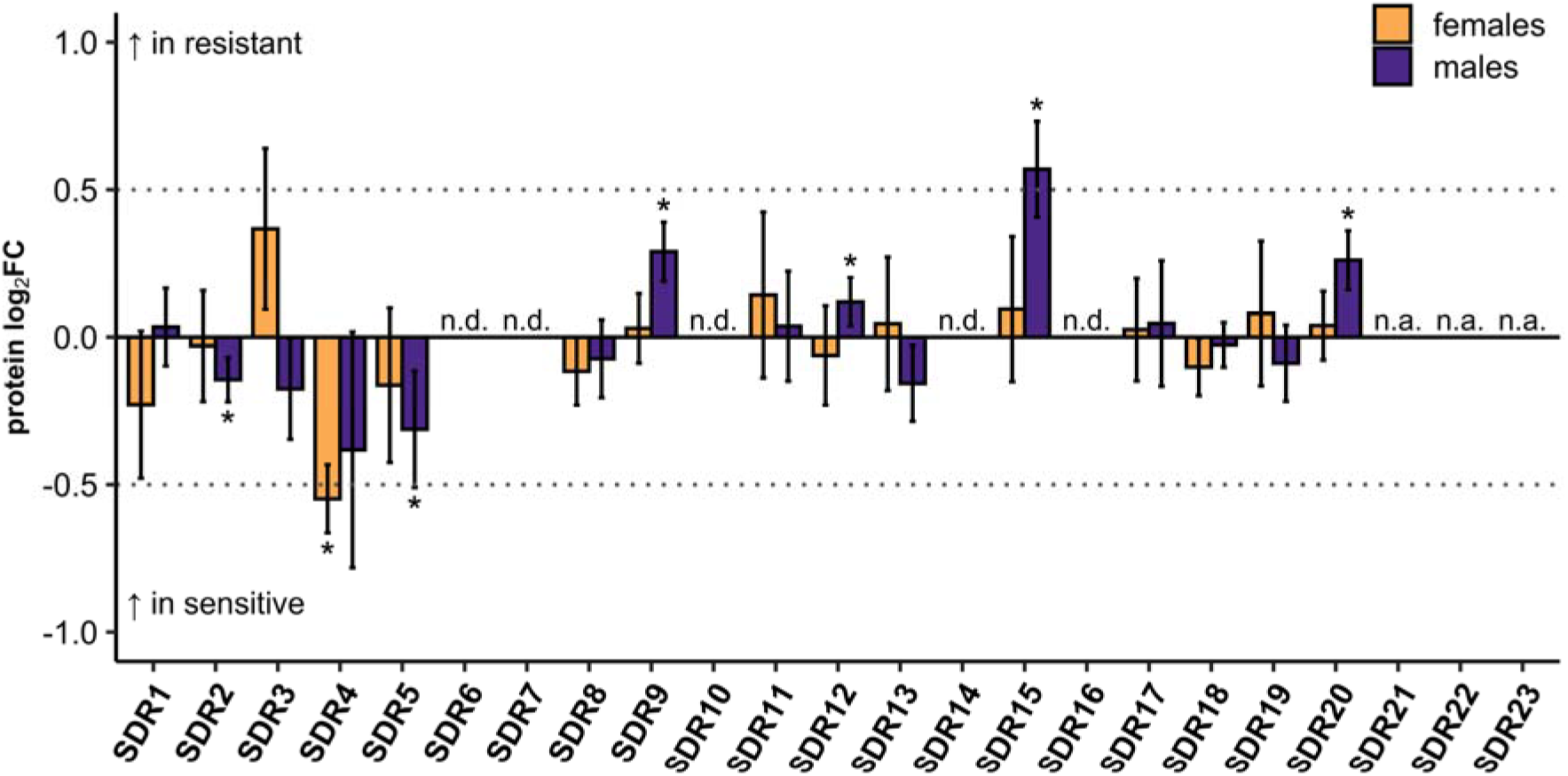
Relative comparison of the constitutive protein expression of SDRs between adults from the drug-sensitive (ISE) and benzimidazole-resistant (IRE) strains. Relative protein expression is displayed as log_2_ fold changes (FC) between adults from the IRE strain compared to the ISE strain. Females and males were compared separately. Protein abundances were determined by targeted MRM/PRM analysis (n = 4). For each peptide, signal intensity of the endogenous light peptide was normalized to that of the corresponding stable isotope labelled heavy peptide. Protein-level values were then calculated from the normalized peptide light-to-heavy ratios. Pairwise group differences were assessed using linear regression models, and statistical significance was determined using Benjamini–Hochberg–adjusted p-values. Error bars represent 95% confidence intervals; asterisks indicate statistically significant differences after multiple testing correction (adjusted *p* < 0.05). Horizontal dotted lines indicate log_2_FC of ± 0.5. Abbreviations: n.d., not detected; n.a., not analyzed

### Effect of FLU on the mRNA and protein expression of SDRs in adults of *H. contortus*

Adults of *H. contortus* of both the ISE and IRE strains were subsequently exposed to FLU (0.01, 1 or 5 µM) for 4 or 12 hours, and the transcriptional response of 23 SDR genes was analyzed.

The results for the ISE and IRE strain are presented in Figure 3 and Figure 4, respectively.

**Figure 3.**
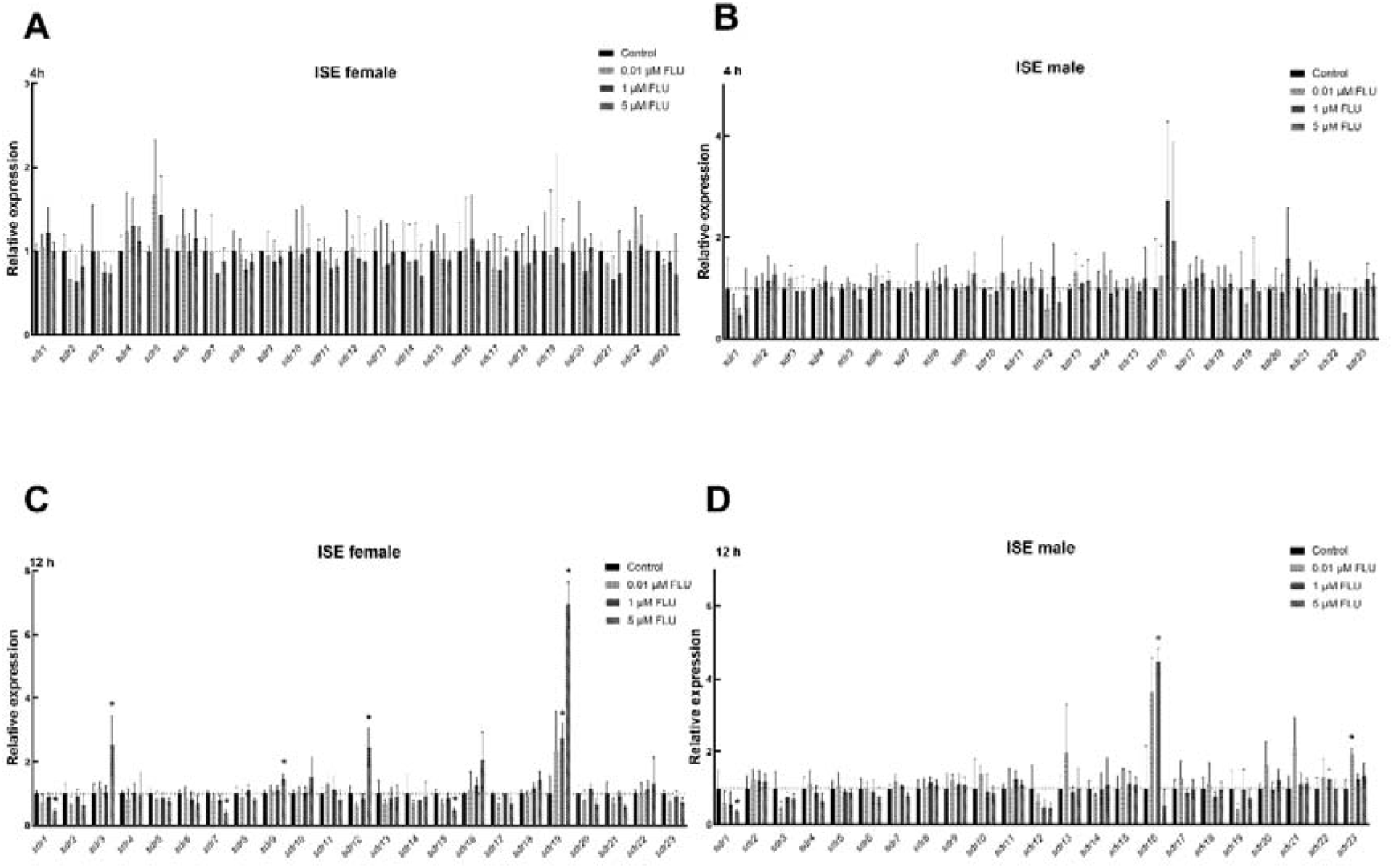
Transcriptional response of SDR genes to flubendazole (FLU) treatment in adult females and males from the drug-sensitive (ISE) strain of *H. contortus*. Relative mRNA expression levels of 23 SDRs were measured in adult females (A, C) and adult males (B, D) after 4 h (A, B) and 12 h (C, D) of incubation. Gene expression is presented as the mean ΔΔCt fold change (n = 2-4). Error bars indicate standard deviation. Asterisks denote significant differences between treated and untreated samples (adjusted *p* < 0.05). Statistical analysis was performed using two-way ANOVA followed by a *post-hoc* Dunnet’s test.

**Figure 4.**
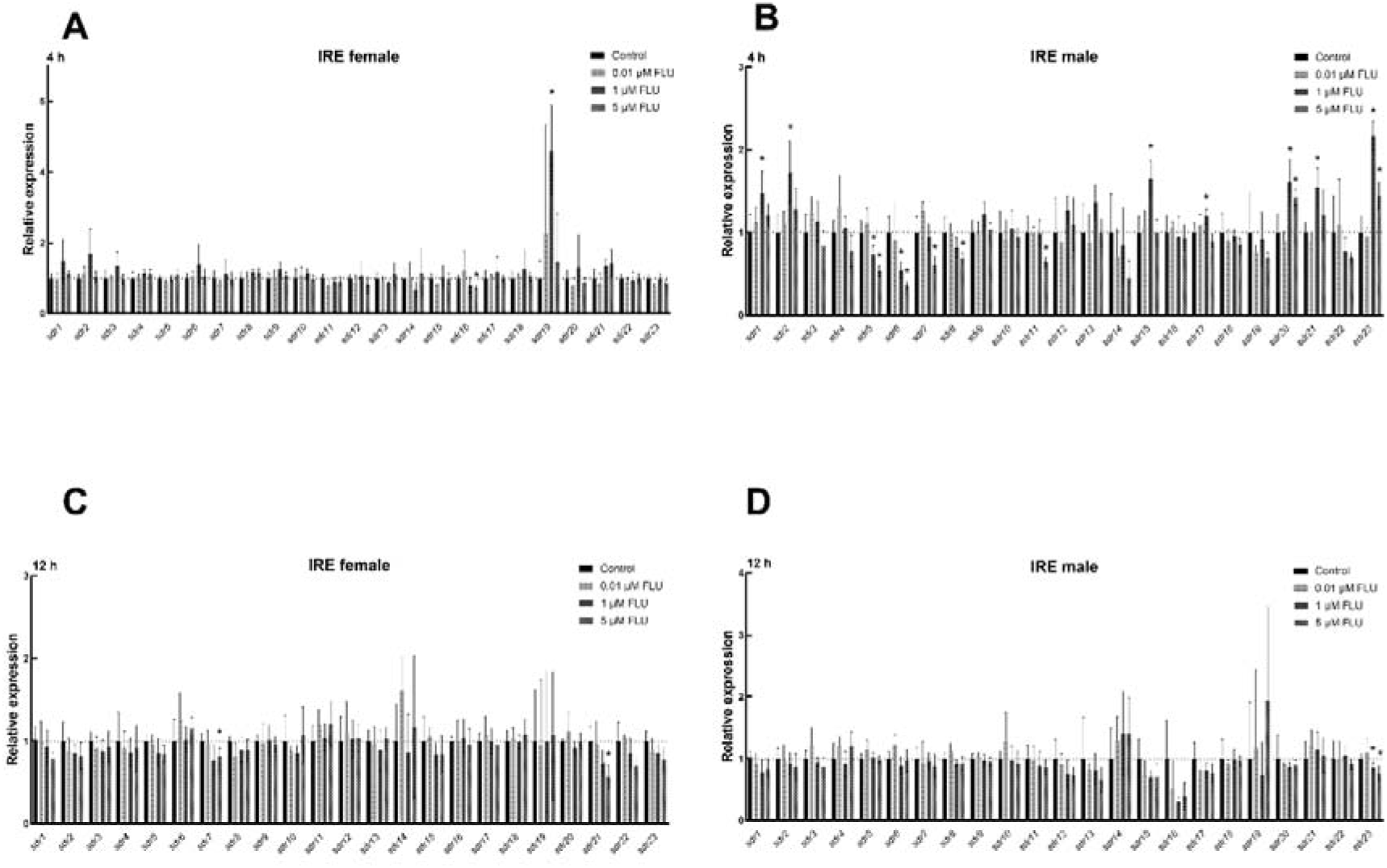
Transcriptional response of SDR genes to the flubendazole (FLU) treatment in female and male adults from the benzimidazole-resistant (IRE) strain of *H. contortus*. Relative mRNA expression levels of 23 SDRs were measured in adult females (A, C) and adult males (B, D) after 4 h (A, B) and 12 h (C, D) of incubation. Gene expression is presented as the mean ΔΔCt fold change (n = 2-4). Error bars indicate standard deviation. Asterisks denote significant differences between treated and untreated samples (adjusted *p* < 0.05). Statistical analysis was performed using two-way ANOVA followed by a *post-hoc* Dunnet’s test.

Overall, the effects of FLU on the mRNA expression of SDRs were concentration-dependent, with most changes observed at the highest tested concentration (5 µM). In addition, the transcriptional responses to FLU were highly sex- and strain-specific, with males from the IRE strain exhibiting the most FLU-induced changes in the SDRs expression. Interestingly, most FLU-mediated alterations were detected after 12-hour exposure. However, in adults from the IRE strain (especially males), the strongest transcriptional response occurred after 4-hour exposure.

The expression of only three SDRs was changed jointly in females or males from both strains, with females exhibiting a decrease in *sdr7* and an increase in *sdr19* levels, while males showed dysregulated expression of *sdr23*. Besides, in females of the ISE strain, FLU increased predominantly the expressions of *sdr3* and *sdr12* but decreased that of sdr15. In males of the ISE strain, FLU exposure led mainly to the increase of sdr16 expression. In females of the IRE strain, a drop in the expression of *sdr21* was the most dominant response to the treatment. And lastly, in males from the IRE strain, several SDRs exhibited decreased (*sdr5*, *sdr6*, *sdr7*, *sdr8*, and *sdr11*) or increased (*sdr1*, *sdr2*, *sdr15*, *sdr20*, *sdr21*, and *sdr23*) expression levels after 4-hour exposure; however, most of these changes did not persist after longer incubation.

At the protein level, a mild but significant response to 24-hour exposure to FLU (5 µM) was observed exclusively in adults of the ISE strain (Figure 5). In females, FLU increased the expression of SDR11 (+28 %) and at the same time decreased the expression of SDR17 (−16 %). On the other hand, males were found to respond with elevated levels of SDR9 (+10 %), SDR19 (+14 %), and SDR20 (+17 %), along with decreased relative amounts of SDR4 (−18 %) and SDR8 (−12 %). Interestingly, the number of significantly dysregulated SDRs was higher in males than females, which is in good agreement with the fact that most of the resistance-related changes in the SDR protein expression were observed in males (Figure 2).

**Figure 5.**
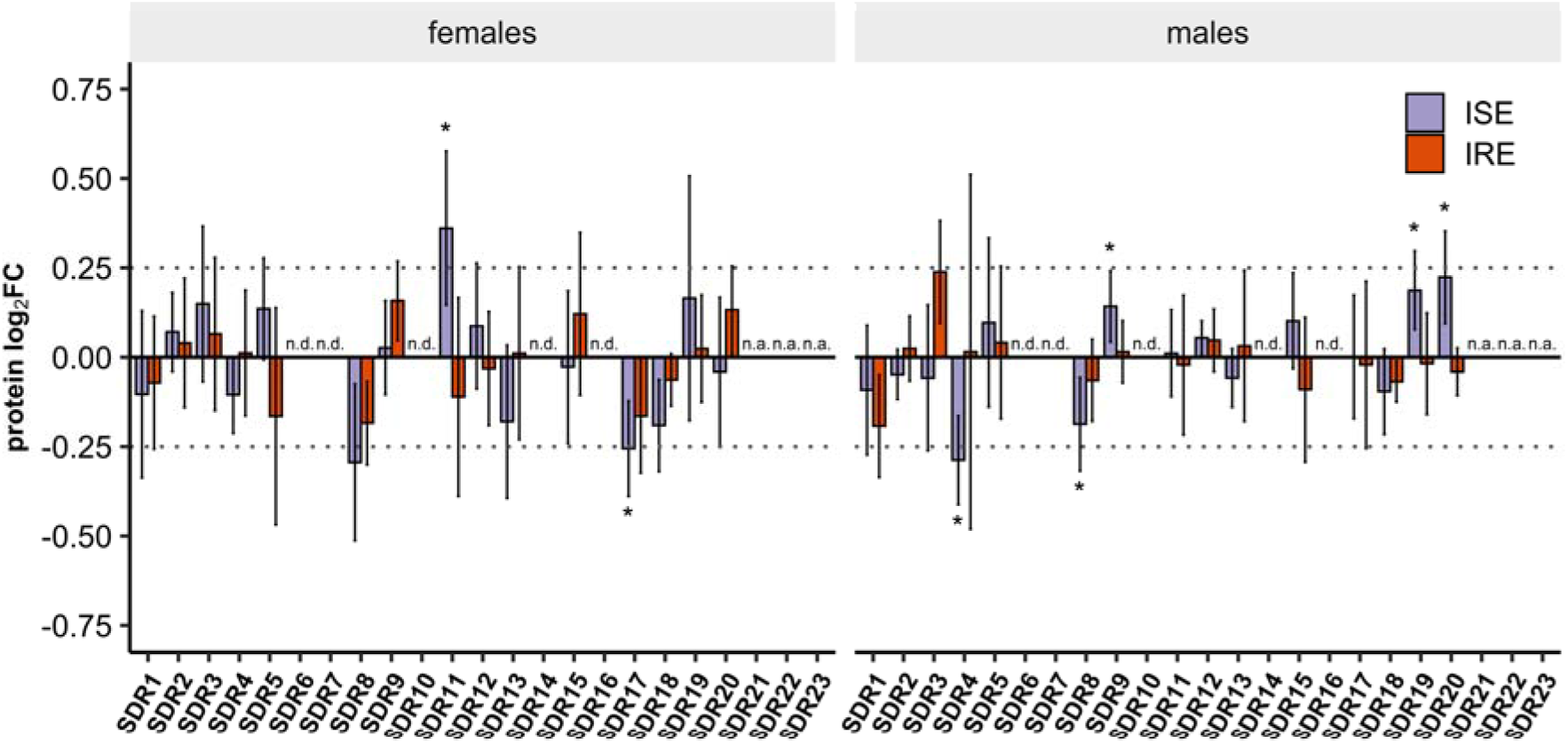
Effect of flubendazole (FLU) on the protein expression of SDRs in the adults of *H. contortus*. Adults were treated with 5 µM FLU for 24 hours. Relative protein expression is displayed as log_2_ fold changes (FC) between treated and untreated adults from the benzimidazole-resistant and sensitive strains separately. Protein abundances were determined by targeted MRM/PRM analysis (n = 4). For each peptide, signal intensity of the endogenous light peptide was normalized to that of the corresponding stable isotope labelled heavy peptide. Protein-level values were then calculated from the normalized peptide light-to-heavy ratios. Pairwise group differences were assessed using linear regression models, and statistical significance was determined using Benjamini–Hochberg–adjusted p-values. Error bars represent 95% confidence intervals; asterisks indicate statistically significant differences after multiple testing correction (adjusted *p* < 0.05). Horizontal dotted lines indicate log_2_FC of ± 0.25. Abbreviations: n.d., not detected; n.a., not analyzed

### Effect of FLU on the mRNA expression of SDRs in juvenile stages of *H. contortus*

Lastly, we investigated whether the treatment with FLU can also influence the mRNA expression of the 23 investigated SDRs in the earlier developmental stages of *H. contortus* (Additional file 6). We found no transcriptional response to the FLU exposure in eggs, L1, or L3 larvae. Only xL3 exposed to 0.01 µM FLU exhibited significantly increased expression of *sdr1*. However, higher FLU concentration (1 µM) did not induce either *sdr1* or other tested SDRs.

## Discussion

SDR superfamily consists of many interesting enzymes given their key roles in signaling pathways, regulation and defense mechanisms [11, 26]. Similarly, the importance of certain SDR members in nematodes has been reported in several studies [27–30]. In *H. contortus*, genome sequencing enabled the identification of 46 members of the SDR family. However, information about these enzymes in this parasitic nematode is still limited, despite the potential of SDRs to serve as drug targets or contribute to drug resistance. To address these knowledge gaps, genome localization, phylogenetic, and transcriptomic analysis of SDRs in *H. contortus* were performed in our previous study [14]. In addition, a functional analysis of SDR12 was performed, revealing its reductase activity towards various substrates, yet no activity towards the reduction of FLU [31]. Building upon these findings, the present study investigated the protein expression of SDRs and their inducibility by the anthelmintic drug FLU. Of note, FLU is inactivated by carbonyl reduction in nematodes [5, 9]; therefore, increased expression or activity of SDRs (or other carbonyl-reducing enzymes) could contribute to enhanced FLU tolerance in resistant nematodes.

Comparison of the constitutive levels of SDRs in females and males revealed significant sex-associated differences in both transcript and protein expression, which is in good agreement with our previously reported findings of pronounced sex-specific differences in the global proteomes of adult *H. contortus* [32]. Notably, the constitutive expression patterns of several SDRs were consistent between both molecular levels investigated, showing a coordinated increase in SDR4, SDR5, and SDR8 in males, and, conversely, a coordinated increase in SDR13, SDR17, SDR18, and SDR20 in females. In the case of two SDRs (SDR2 and SDR11), their sex-related protein abundance was discordant with their mRNA expression, which might indicate involvement of posttranscriptional regulation in their expression.

Resistance-associated changes in the protein levels of the investigated SDRs were less pronounced than the sex-related changes discussed above. This observation is again consistent with our previous findings, in which drug resistance exerted a considerably weaker effect on the overall proteomic profiles than parasite sex [32]. In addition, most resistance-related differences were observed in males, with four SDR isozymes (SDR9, SDR12, SDR15, and SDR20) exhibiting increased abundances in benzimidazole-resistant males. A similar trend was observed in our previous work, where resistance-associated changes in the mRNA levels of multiple SDR genes occurred exclusively in males [14]. However, we found that the transcriptional alterations are not consistently mirrored at the protein level. Specifically, SDR9, SDR12, SDR15, and SDR20 were significantly upregulated at the protein level, whereas only SDR12 also showed significantly upregulation at the mRNA level. From a pharmacological perspective, enzyme abundance is more relevant than mRNA expression, and all SDRs with elevated protein levels in the benzimidazole-resistant strain warrant further investigation regarding their potential role in drug resistance.

To better assess the significance of SDRs as potential participants in FLU inactivation and contributors to drug resistance in *H. contortus*, it is also necessary to investigate their inducibility by this drug. In fact, the inducibility of SDRs by various xenobiotics has been well-documented not only in humans [33], but also in nematodes. In *C. elegans*, the exposure to a common persistent pollutant 2,2’,5,5-tetrachlorobiphenyl and other xenobiotics caused an increased expression of several SDR enzymes [34, 35]. In pine wood nematode (*Bursaphelenchus xylophilus),* several SDRs were among the genes upregulated by β-pinene-mediated stress [36]. Similarly, anthelmintic drugs ivermectin, pyrantel, and thiabendazole induced the expression of several SDRs in *Parascaris univalens*, a pathogenic parasite of foals and yearlings [37]. On the other hand, FLU induced the mRNA expression of another class of reductases in adults of *H. contortus* – aldo/keto reductases, specifically three genes designated as *akr19*, *akr9*, and *akr15* [38]. In addition, the ability of FLU to increase the activity of carbonyl reducing enzymes was observed in sheep and pheasant [39, 40]. However, the mechanism of FLU-mediated induction of these enzymes is still unknown.

In our study, the transcriptional response to exposure to sublethal concentrations of FLU was tested across all developmental stages of *H. contortus*. Given that more than 50% of administered FLU is excreted unchanged in the faeces of treated animals [6], free-living stages of helminths on pastures may be also affected by environmental exposure to the drug through contaminated fecal matter. However, we observed almost no effect of FLU on the expression of SDRs in the juvenile stages of *H. contortus*.

In adults, on the other hand, the contact with FLU resulted in evident sex- and strain-specific changes in both the mRNA and protein expression of several SDRs. Interestingly, adults from the benzimidazole-resistant strain (particularly males) exhibited faster and stronger transcriptional response to FLU than adults from the sensitive strain; however, FLU-mediated changes in the protein abundance were observed exclusively in the sensitive strain (particularly in males). Surprisingly, there was no agreement between the significant expression changes identified at the two levels. This discrepancy in the response may arise from several factors: (i) while 23 SDRs were quantified at the mRNA level, only 15 SDRs could be analyzed at the protein level; (ii) selected duration of incubations (4 and 12 hours for mRNA expression analysis, 24 hours for targeted proteomic analysis) might be too short or too long to observe concordant changes in expression; (iii) other transcriptional and (post)translational mechanisms in the regulation of SDRs might be involved that could not be addressed by our experimental design. Despite these experimental limitations, our results clearly demonstrate the inducibility of some SDRs in *H. contortus* by exposure to FLU at sublethal doses.

It is important to mention that males rather than females of *H. contortus* consistently exhibited much more pronounced resistance-related alterations in the mRNA [14] and protein expression of SDRs and simultaneously produced faster and stronger transcriptional and protein-level response to FLU treatment. Two SDRs – SDR9 and SDR20, which protein levels were found elevated after the treatment with FLU in the sensitive males, were also among the handful of SDR enzymes upregulated in the benzimidazole-resistant males. In addition, two other FLU-inducible isozymes SDR16 and SDR23 show higher constitutive expression in all juvenile stages of the IRE strain and in IRE males, respectively [14]. The contribution of these four isozymes to drug resistance and increased elimination of benzimidazole anthelmintics in *H. contortus* warrants further study.

In conclusion, the constitutive mRNA and protein expression of the 23 investigated SDRs differed significantly between males and females from both the drug-sensitive and the benzimidazole-resistant strains of *H. contortus*, which highlights the necessity to perform all experiments in both sexes separately. The resistance-associated adaptations were overall less pronounced; however, SDR9, SDR12, SDR15, and SDR20 exhibited higher protein abundances in benzimidazole-resistant males. Juvenile stages of *H. contortus* and adult females did not respond to exposure to sublethal doses of FLU. Conversely, several SDRs were found to be inducible by FLU in both sensitive and benzimidazole-resistant males. Among these, SDR9, and SDR20 are of particular interest, as they not only respond to exposure to the anthelmintic but also exhibit elevated constitutive expression in the resistant strain.

## Data availability

The mass spectrometry proteomics data have been deposited to the ProteomeXchange Consortium (http://proteomecentral.proteomexchange.org) via the Panorama Public [41] partner repository with the dataset identifier PXD081312. Skyline documents containing the targeted mass spectrometry data can be accessed on the Panorama Public website https://panoramaweb.org/Hco_SDRs.url). The experimental metadata has been generated using lesSDRF [42]. Further datasets and results supporting the conclusions of this article are included within the additional files.

## Acknowledgement

We would like to thank Prof. Marián Várady of the Institute of Parasitology, Slovak Academy of Sciences for providing *H. contortus* larvae.

## Authors contributions

Conceptualization: LS, LRS, PM; Methodology: LRS, MŠ, KŠ, PM, NS, TR, ML; Software: MŠ, PM, NS, NL; Validation: MŠ, NR, PM; Formal analysis: MŠ, PM; Investigation: MŠ, KŠ, PM, NS; Resources: LRS, PM, TR; Data curation: MŠ, KŠ, PM; Writing – Original Draft: LS, NS, KŠ, MŠ; Writing – Review & Editing: PM, LRS, MŠ, NL, ML; Visualization MŠ, PM; Supervision: LS, LRS, PM, TR, ML; Project administration: LRS, PM; Funding acquisition: LRS, PM, TR

## Declarations

The authors declare no conflict of interest.

## Ethics approval and consent to participate

All experimental procedures involving sheep were evaluated and approved by the Ethics Committee of the Ministry of Education, Youth and Sports of the Czech Republic (project no. MŠMT-20144/2023-4).

